# Neuropeptide Y regulation of L-type Ca^2+^ channel activity is altered following chronic myocardial infarction

**DOI:** 10.1101/2025.11.25.690298

**Authors:** Chase M. Fiore, Shailesh R. Agarwal, Seyi E. Elasoru, Jeffrey Ardell, Olujimi Ajijola, Kalyanam Shivkumar, Robert D. Harvey

**Affiliations:** Department of Pharmacology, University of Nevada, Reno, Reno, NV; Cardiac Arrhythmia Center, UCLA, Los Angeles, CA

## Abstract

Neuropeptide Y (NPY) is a co-transmitter released from sympathetic neurons along with norepinephrine (NE). It has been observed that cardiac NPY levels are significantly elevated following myocardial infarction (MI), and this has been linked to an increase in ventricular arrhythmogenicity associated with elevated sympathetic tone.

However, the effects that NPY has on the electrical activity of ventricular myocytes remain poorly understood. Previous studies have examined the influence of NPY alone on cardiac ion channel function, but not in the presence of NE, which is the situation expected in vivo. Furthermore, no one has examined the effects of NPY on ion channel activity following MI. The present study explored the impact of NPY on the L-type Ca^2+^ current in ventricular myocytes isolated from the hearts of normal healthy pigs and pigs subjected to MI. We found that NPY alone has a stimulatory effect on the Ca^2+^ current in myocytes isolated from healthy pigs. However, in the presence of NE, the effect of NPY was inhibitory. The stimulatory effect of NPY alone was blocked by the Y_1_ receptor antagonist BIBO3304, while the inhibitory effect observed in the presence of NE was blocked by the Y_2_ receptor antagonist BIIE0246. When the effects of NPY were examined using hearts from pigs following recovery from MI, the stimulatory effect of NPY was absent in myocytes obtained from both remote and border zone areas of infarcted hearts. The inhibitory effect of NPY observed in the presence of NE was also absent in myocytes from remote areas of the infarcted heart. However, the inhibitory effect of NPY observed in the presence of NE was intact in border zone cells. The implications of these results are discussed as they relate to the potential arrhythmogenic effects of NPY following MI.

**Graphical Abstract. NPY exerts bimodular, context dependent effects on LTCC.:** 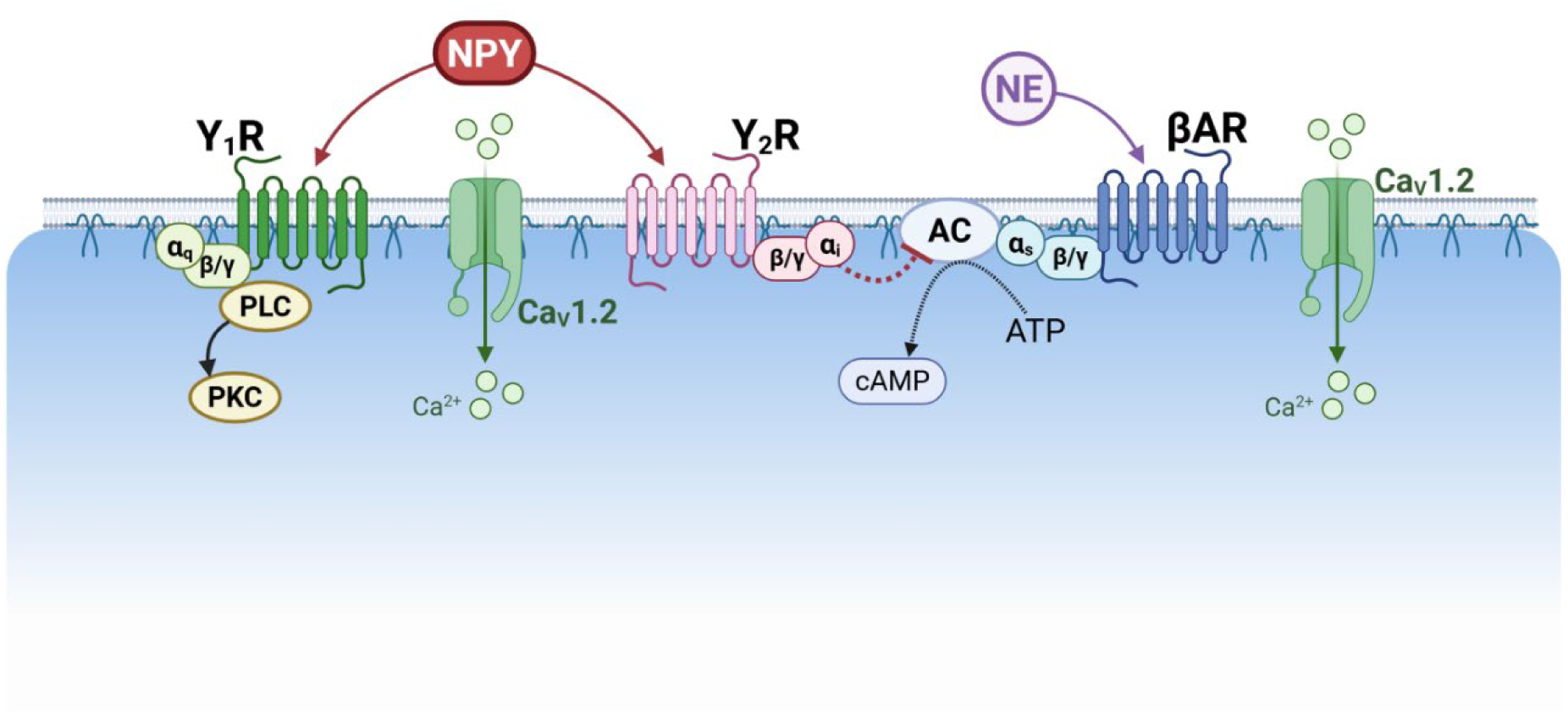

Proposed signaling pathways for diverse effects of NPY in ventricular cardiac myocytes. Norepinephrine (NE) and neuropeptide Y (NPY) co-application results in a Y_2_/G_i_-mediated reduction in β-adrenergic (βAR)/G_s_ enhanced I_CaL_, while NPY alone modestly enhances the current via Y_1_/G_q_ mechanism. These pathways are regionally altered following chronic MI.

## Introduction

The sympathetic nervous system plays an essential role in increasing cardiac output in response to stress and exercise by regulating both heart rate and contractility. These effects are most commonly associated with the release of the neurotransmitter norepinephrine (NE) and subsequent β-adrenergic receptor (βAR) activation. However, following myocardial infarction (MI), sympathetic stimulation also produces maladaptive responses that contribute to the development of heart failure [1,2]. Furthermore, the elevation in sympathetic tone that occurs during heart failure is associated with an increased incidence of ventricular arrhythmias leading to sudden cardiac death (SCD) [2–4]. Consistent with this fact, βAR antagonists are the only class of antiarrhythmic drugs found to reduce the frequency of arrhythmias and SCD in heart failure patients [5–7]. Despite the use of β-blockers in this situation, in the United States more than 300,000 people a year still succumb to SCD [8]. This suggests that other factors, in addition to βAR stimulation, contribute to sympathetic induced arrhythmias.

Another potential element contributing to arrhythmogenesis is neuropeptide Y (NPY). This 36 amino acid peptide is highly expressed in sympathetic neurons, and it is released as a co-transmitter along with NE, especially when sympathetic tone is high [9–11]. The idea that this neuropeptide may contribute to arrhythmogenesis associated with SCD is supported by the finding that coronary sinus levels of NPY are elevated following MI, and elevated NPY levels correlate with increased mortality in heart failure patients [12]. In addition, blocking the effects of NPY antagonists has been shown to reduce the threshold for triggering ventricular arrhythmias following MI [13].

Despite evidence that NPY may be arrhythmogenic, the mechanism involved is unknown. One possible explanation is that NPY enhances L-type Ca^2+^ channel activity, which is also a factor contributing to βAR induced arrhythmias [14]. While a few studies have looked at the effects of NPY on L-type Ca^2+^ channel activity, the reported results have been inconsistent [15–17]. Furthermore, all studies to date have looked at responses to NPY alone in myocytes obtained from normal hearts. No experiments have looked at the effects of NPY in the presence of NE, which is the situation that occurs *in situ*. Furthermore, no one has looked at the effects of NPY, either alone or in the presence of NE, in myocytes obtained from hearts that have been subjected to MI. In the present study, we examined the effects of NPY on the L-type Ca^2+^ current recorded from ventricular myocytes isolated from the hearts of normal pigs and pigs subjected to MI. Furthermore, we examined the response to NPY alone as well as the response to NPY in the presence of NE.

## Materials and Methods

### Cell Isolation

Experiments were conducted using cardiac myocytes isolated from the left ventricle of hearts obtained from Yucatan mini pigs (N = 18; 34 ± 0.7 kg). Animals were sedated with intramuscular injection of ketamine (100 mg), intubated, and then anesthetized using isoflurane gas (2 to 4%) inhalation. The heart was then removed following thoracotomy and rinsed free of blood in ice cold cardioplegia solution containing (in mM): NaCl 130, KCl 27, MgSO_4_ 1.2, KH_2_PO_4_ 1.2, glucose 50, and HEPES 6 (pH 7.2). A section of the left ventricular free wall (∼1.5 in x 1.5 in) containing a major coronary artery was attached to a Langendorff apparatus. The tissue was perfused at a constant pressure with Ca^2+^-free physiologic saline solution (PSS) containing (in mM): NaCl 140, KCl 5.4, MgCl_2_ 2.5, glucose 11, HEPES 5.5 (pH 7.4) at 37°C. Coronary artery branches were tied off to maintain perfusion pressure. After 5 to 10 min, the solution was switched to Ca^2+^-free PSS containing collagenase (Worthington Biochemical type II, ∼200 units/ml) and protease (Sigma type XIV, ∼0.1 mg/ml). After 15 minutes of digestion, the tissue was placed in KB solution containing (in mM): K^+^-glutamate 110, KH_2_PO_4_ 10, KCl 25, MgSO_4_ 2, taurine 20, creatine 5, EGTA 0.5, glucose 20, HEPES 5 (pH 7.4). Softened tissue was cut into small pieces, minced, triturated with a blunt transfer pipette, and then filtered through 300 μm^2^ nylon mesh. Cells were rinsed twice in KB solution, before resuspending the pellet in normal Ca^2+^ containing solution.

### Infarct Model

Experiments looking at the effects of chronic myocardial infarction (MI) were conducted using myocytes isolated from the hearts of pigs in which an antero-apical infarct was created by microbead occlusion of the left anterior descending coronary artery at the second diagonal branch via percutaneous catheter, as described previously [3]. The animals were then allowed to recover for 4 to 6 weeks, during which time they were monitored by echocardiography. After verifying that left ventricular ejection fraction was reduced and wall motion was altered, the hearts were removed and myocytes isolated using the method described above. The infarct area was easily identified by visual inspection. After enzymatic digestion, myocytes were isolated from either the infarct border zone or an area remote from the infarct.

The protocols used are in accordance with the *Guide for the Care and Use of Laboratory Animals* as adopted by the National Institutes of Health and approved by the Institutional Animal Care and Use Committees at the University of Nevada, Reno (normal hearts) or the University of California Los Angeles (MI hearts). In all cases, cells were used for experiments on the day of isolation.

### Data Acquisition

Membrane currents were recorded from isolated myocytes using the whole-cell configuration of the patch-clamp technique. All experiments were performed at 37°C using borosilicate glass electrodes polished to a resistance of 1–2 MΩ. Cells were dialyzed with a pipette solution containing (in mM): CsCl 130, TEA-Cl 20, EGTA 5, Mg-ATP 5, and Tris-GTP 0.06, and HEPES 5.0 (pH 7.2). Cells were bathed in an extracellular solution containing (in mM): NaCl 137, CsCl 5.4, MgCl₂ 0.5, CaCl₂ 1.0, NaH₂PO₄ 0.33, glucose 5.5, and HEPES 5.0 (pH 7.4). These solutions set the Cl^-^equilibrium potential at 0 mV.

The L-type Ca^2+^ current was recorded using a MultiClamp 700B amplifier, Digidata 1440 digitizer and pCLAMP 11 software (Molecular Devices). Currents were lowpass filtered at 4 kHz and sampled at 10 kHz. Membrane potential was held at -80 mV. A 50 ms pre-pulse to -40 mV was used to inactivate Na^+^ channels. This was followed by a 100 ms test pulse to 0 mV to elicit the L-type Ca^2+^ current. Because pig ventricular myocytes were found to express the cAMP regulated Cl^-^ current, the effect that changes in this time-independent background current might have on measurement of Ca^2+^ current responses was eliminated by recording the Ca^2+^ current at the Cl^-^equilibrium potential (0 mV).

The time course of changes in the amplitude of the Ca^2+^ current was monitored by recording the amplitude of the peak inward current elicited during the test pulse to 0 mV applied once every 5 s. After establishing a stable baseline, cells were exposed to external solutions containing various drugs, as indicated. Norepinephrine (NE), neuropeptide Y (NPY), BIBO3304 (BIBO), and BIIE0246 (BIIE), were all purchased from Tocris Biosciences. Solutions containing NE were made from aqueous stock solutions prepared fresh daily. The remaining drug containing solutions were made fresh daily from stock solutions prepared in DMSO and stored at -20°C.

### Data Analysis

Average responses (mean ± SEM) were determined by measuring drug effects in the number of myocytes indicated (n) obtained from the number of animals indicated (N). Statistical analysis was by paired or unpaired students t-test, with correction for multiple comparisons where needed, using Prism software (GraphPad version 10.4.2).

## Results

Experiments were conducted using ventricular myocytes isolated from the hearts of the Yucatan minipig. This model was chosen because its anatomical and electrophysiological properties more closely mimic those of the human heart. This makes it an ideal large animal model for translational studies investigating mechanisms of cardiac arrhythmias, especially those involving MI [18,19].

### Effects of NPY in Control Myocytes

Previous studies have reported that NPY alone either stimulates, inhibits or has no effect on the L-type Ca^2+^ current in ventricular myocytes from other animal models [15–17]. Therefore, we first sought to establish the effect that NPY has on the Ca^2+^ current in ventricular myocytes from normal healthy pigs (Figure 1A-C). It was found that exposure to 100 nM NPY enhanced the baseline current density from 5.0 ± 0.57 pA/pF to 6.4 ± 0.48 pA/pF, which represents an increase of 15.4 ± 2.76% (p = 0.0002, N/n = 3/13; Figure 1C).

**Figure 1.**
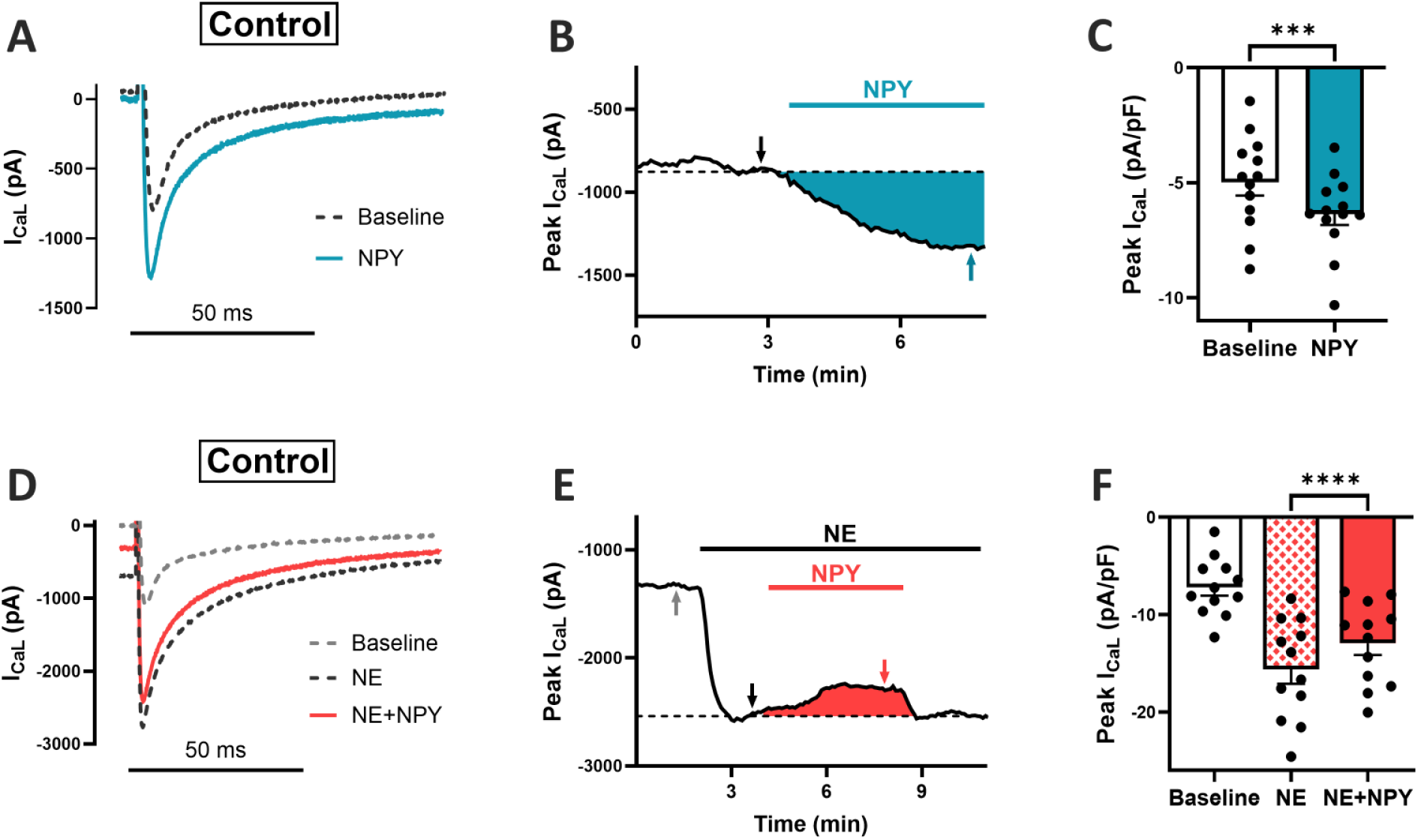
Neuropeptide Y (NPY) exerts bimodal regulation of I_CaL_ in control PVM. (A) Representative whole-cell I_CaL_ traces recorded under baseline conditions (dashed black line) and following application of 100 nM NPY (blue line). (B) Time course highlighting the NPY-induced increase in peak I_CaL_ amplitude from an example myocyte. (C) Aggregate data showing a significant increase of 15.4 ± 2.76% in peak I_CaL_ following NPY application (baseline: –4.99 ± 0.57 pA/pF; NPY: –6.36 ± 0.48 pA/pF; p = 0.0002, N/n = 3/13). (D) Representative I_CaL_ traces recorded under baseline conditions (dashed gray line), with 30 nM norepinephrine (NE; dashed black line) and following co-application of 100 nM NPY (red line). (E) Time course illustrating suppression of I_CaL_ amplitude by NPY during adrenergic stimulation. (F) Pooled data reveal a significant reduction (32.7 ± 4.23%) of peak current elicited by NE upon addition of NPY (NE: – 15.6 ± 1.48 pA/pF; NE+NPY: –12.9 ± 1.22 pA/pF; p < 0.0001, N/n = 4/12).

Sympathetic neurons innervating the heart have been shown to release NPY together with NE [10]. Therefore, to assess the effect of NPY under more physiological conditions, we measured the Ca^2+^ current response to NPY following exposure to NE (Figure 1D-F). Application of 30 nM NE significantly enhanced the Ca^2+^ current, increasing the peak current density from 7.2 ± 0.85 pA/pF to 15.6 ± 1.48 pA/pF, which represents a 116% increase over baseline (p < 0.0001). This is consistent with the ability of NE to activate β-adrenergic receptors, stimulating adenylyl cyclase (AC), production of cAMP, and subsequent activation of protein kinase A (PKA). Interestingly, rather than observing an additional increase (as might be expected, given the effect of NPY alone), addition of 100 nM NPY in the continued presence of NE reduced the Ca^2+^ current density to −12.9 ± 1.22 pA/pF. This represents a 32.7 ± 4.23% decrease in the response to NE (p < 0.0001, N/n = 4/12; Figure 1F). This data demonstrates a previously unappreciated, modulatory role of NPY on the L-type Ca^2+^ current in cardiac myocytes.

### NPY Receptors Involved

We then attempted to identify the specific type of receptor involved in mediating the effects of NPY. Cardiac myocytes have been reported to express both NPY Y_1_ and Y_2_ receptor subtypes [11]. Y_1_ receptors are associated with activation of G_q_ and protein kinase C (PKC) dependent signaling pathways, while Y_2_ receptors are linked to G_i_, which can inhibit AC activity. PKC has been reported to stimulate L-type Ca^2+^ channel activity in some cardiac myocytes [20], and G_i_ inhibition of AC can antagonize β-adrenergic production of cAMP. Therefore, we tested the hypothesis that the stimulatory effect observed in the absence of NE involves Y_1_ receptors, while the inhibitor effect observed in the presence of NE involves Y_2_ receptors (see Graphical Abstract).

As predicted, exposure to NPY in the presence of the Y_1_ receptor antagonist BIBO3304 (BIBO, 1 μM) blocked the stimulatory effect of NPY alone (Figure 2A-C). In the presence of BIBO alone, the peak Ca^2+^ current density was 7.0 ± 0.78 pA/pF. This was not significantly different from the current density of 7.1 ± 0.66 pA/pF observed following addition of NPY application (p = 0.84, N/n = 2/8; Figure 2C). In contrast, a stimulatory effect of NPY could still be observed. In the presence of BIIE0246 (BIIE, 1 μM) (Figure 2D-F). In the presence of BIIE alone, the peak Ca^2+^ current density was 9.1 ± 1.1 pA/pF. However, following addition of NPY, this increased to 10 ± 1.2. This represents a 10.9 ± 2.60% increase (p = 0.0004, N/n = 3/11; Figure 2F). These findings support the conclusion that the stimulatory effect of NPY in the absence of NE is most likely due to the activation of Y_1_ receptors.

**Figure 2.**
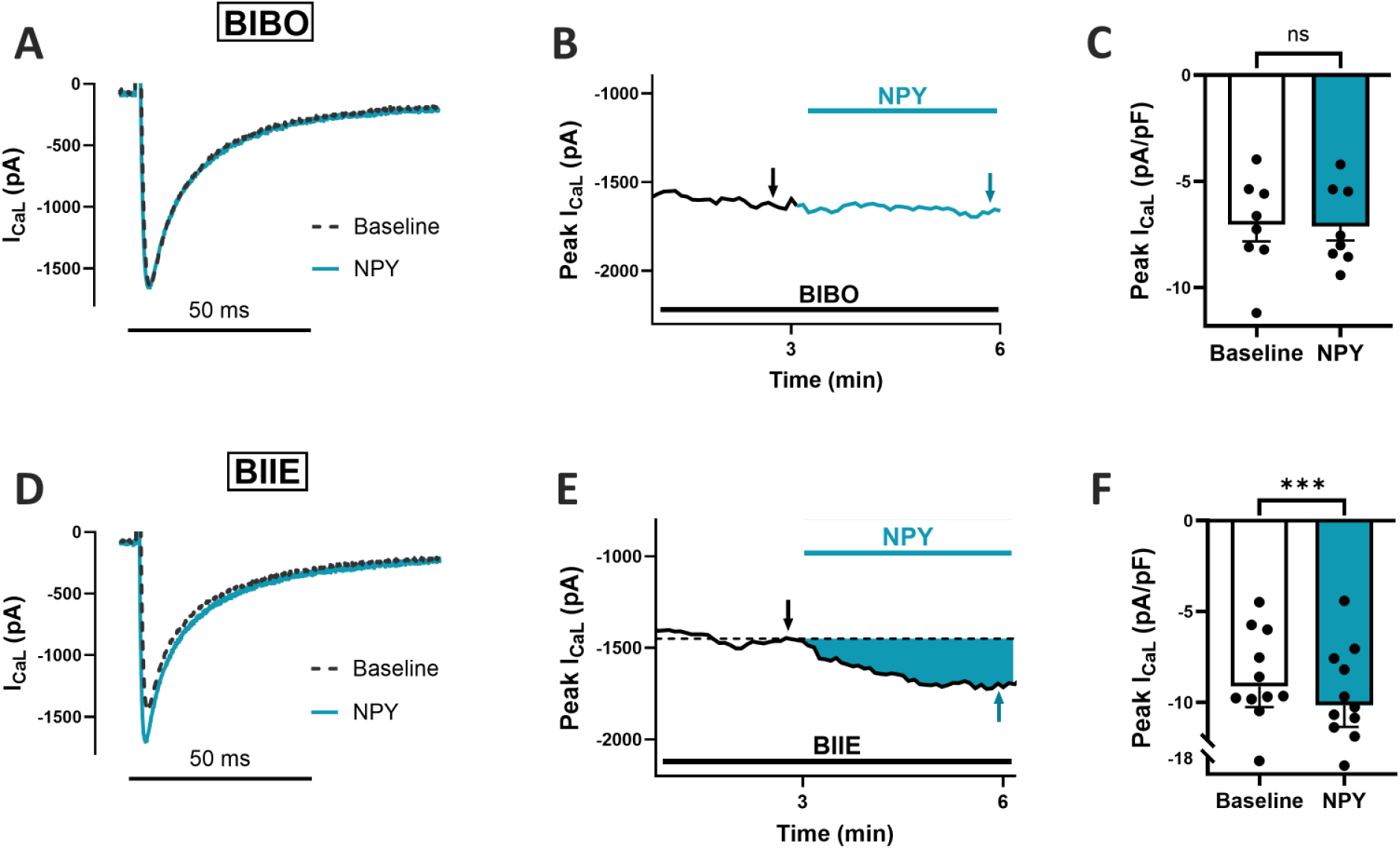
NPY enhancement of I_CaL_ due to Y_1_R activation in control PVM. (A) Representative whole-cell I_CaL_ traces recorded under baseline conditions in the presence of the Y_1_ antagonist BIBO3304 (dashed black line) and following 100 nM NPY application (blue line). (B) Time course showing minimal change (0.21 ± 4.26% increase) in I_CaL_ amplitude during NPY exposure. (C) Aggregate data reveal no significant difference in current density upon administration of NPY (baseline: –7.03 ± 0.78 pA/pF; NPY: –7.12 ± 0.66 pA/pF; p = 0.8365, N/n = 2/8). (D) Representative I_CaL_ traces recorded under baseline conditions in the presence of 1 µM BIIE0246 (dashed black line) and following 100 nM NPY application (blue line). (E) Time course showing a sustained increase in I_CaL_ amplitude during NPY treatment. (F) Summary data show a significant increase of 10.9 ± 2.60% in peak current following addition of NPY (baseline: –9.13 ± 1.13 pA/pF; NPY: –10.2 ± 1.18 pA/pF; p = 0.0004, N/n = 3/11).

We then conducted similar experiments to identify the receptor subtype responsible for the inhibitory effect of NPY observed in the presence of NE. When we repeated experiments like those illustrated in Figure 1D, but in the presence of the Y_1_ receptor antagonist BIBO (Figure 3A-C), the inhibitory effect of NPY remained intact. In the presence of BIBO alone, the peak Ca^2+^ current density was 9.2 ± 0.94%. Following exposure to 30 nM NE, this increased to 24 ± 1.9 pA/pF, which represents a 157% increase over baseline (p < 0.0001). Furthermore, subsequent addition of 100 nM NPY decreased the peak current density to 22 ± 1.6 pA/pF, which represents a 11% decrease in the response to NE (p = 0.011, N/n = 4/9; Figure 3C). In contrast, the inhibitor effect of NPY was blocked in the presence of the Y_2_ receptor antagonist BIIE (Figure 3D-F). In the presence of BIIE alone, the peak Ca^2+^ current density was 8.7 ± 1.2 pA/pF. Following exposure to 30 nM NE, this increased to 21.4 ± 2.40 pA/pF, which represents a 147% increase over baseline (p < 0.0001). However, following subsequent addition of NPY, the peak current density was not significantly affected. It remained at 21.0 ± 2.03 pA/pF (p = 0.56, N/n = 2/10; Figure 3F). These results support the conclusion that the inhibitory effect of NPY observed in the presence of NE is most likely mediated by Y_2_ receptors.

**Figure 3.**
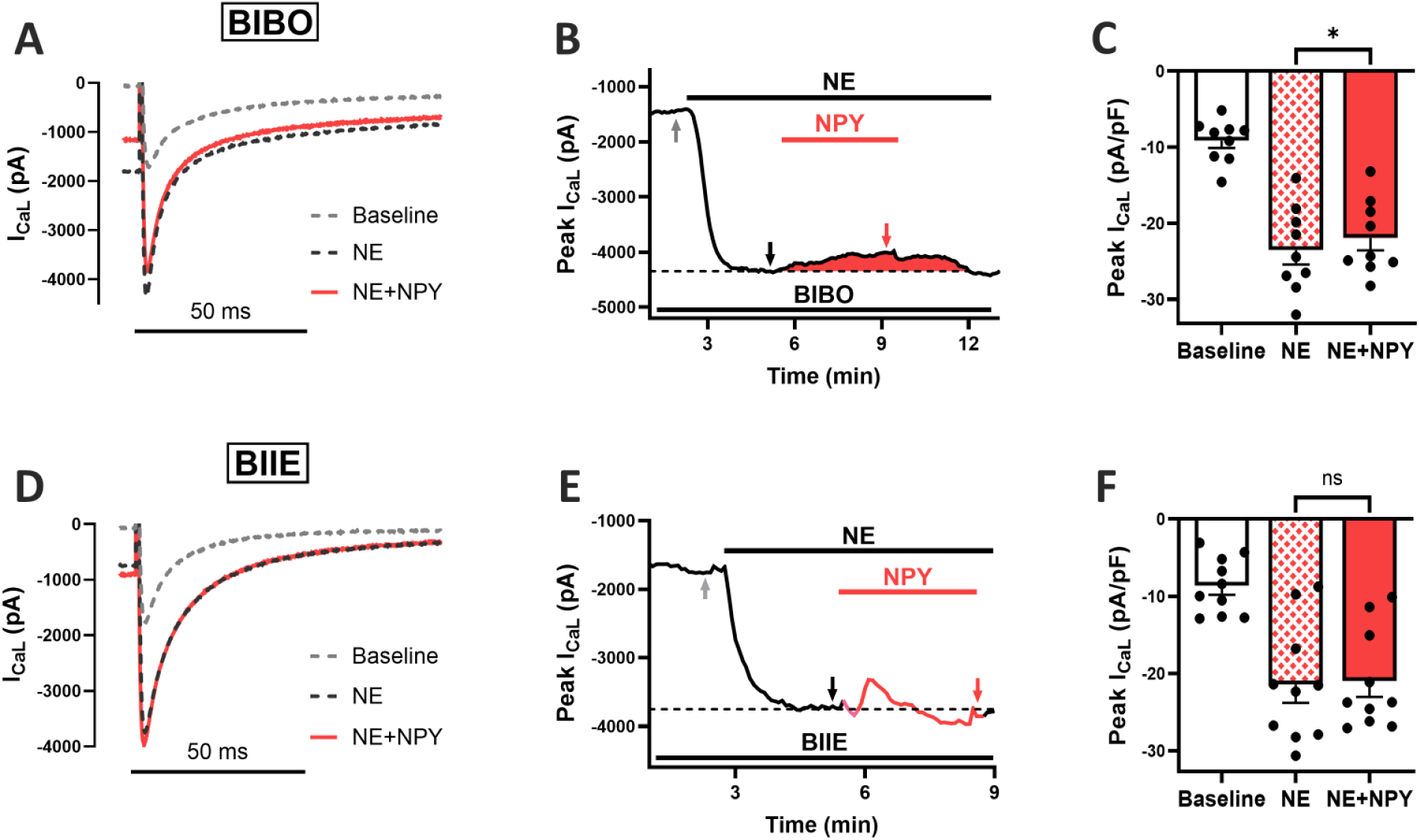
Y_2_R activation is responsible for NPY inhibition of NE-stimulated I_CaL_ in control PVM. (A) I_CaL_ traces recorded under baseline conditions (dashed gray line), following 30 nM NE + 1 µM BIBO3304 (dashed black line) and after NE + 100 nM NPY co-application (red line). (B) Time course demonstrating NPY-mediated reduction of I_CaL_. (C) Average NE stimulated I_CaL_ density was significantly reduced by 11.0 ± 3.04% in the presence of NPY (NE: –23.5 ± 1.88 pA/pF; NE+NPY: –21.9 ± 1.64 pA/pF; p = 0.0106, N/n = 4/9). (D) Representative I_CaL_ traces recorded under baseline conditions (dashed gray line), with 30 nM NE + 1 µM BIIE0246 (dashed black line) and following co-application of NE + 100 nM NPY (red line). (E) Time course showing no sustained suppression of I_CaL_ by NPY. (F) Quantification of steady-state responses shows no significant change (1.06 ± 6.25% decrease) in current amplitude (NE: –21.4 ± 2.40 pA/pF; NE+NPY: –21.0 ± 2.03 pA/pF; p = 0.5631, N/n = 2/10).

### Effects of NPY Following Myocardial Infarction

Next, we determined whether the responses to NPY were altered following MI. These experiments were conducted in two different groups of cells. The first population of cells was isolated from the base of the left ventricle, which was remote from the area of the infarct. The second population of cells was isolated from the infarct border zone. When we examined the response of the Ca^2+^ current to NPY alone, the stimulatory effect that NPY produced in myocytes from healthy hearts was absent in both the remote and border zone cells. In remote zone myocytes (Figure 4A-C), the peak Ca^2+^ current density observed under baseline conditions was 4.6 ± 0.30 pA/pF, and this was not significantly changed following exposure to NPY, where the current density was 4.5 ± 0.28 pA/pF (p = 0.44, N/n = 4/10; Figure 4C). A similar observation was made in border zone myocytes (Figure 4D-F). The peak Ca^2+^ current density was 5.3 ± 0.34 pA/pF under baseline conditions, and this was not significantly altered following exposure to NPY, where the current density was 5.0 ± 0.33 pA/pF (p = 0.1105, N/n = 4/6; Figure 4F).

**Figure 4.**
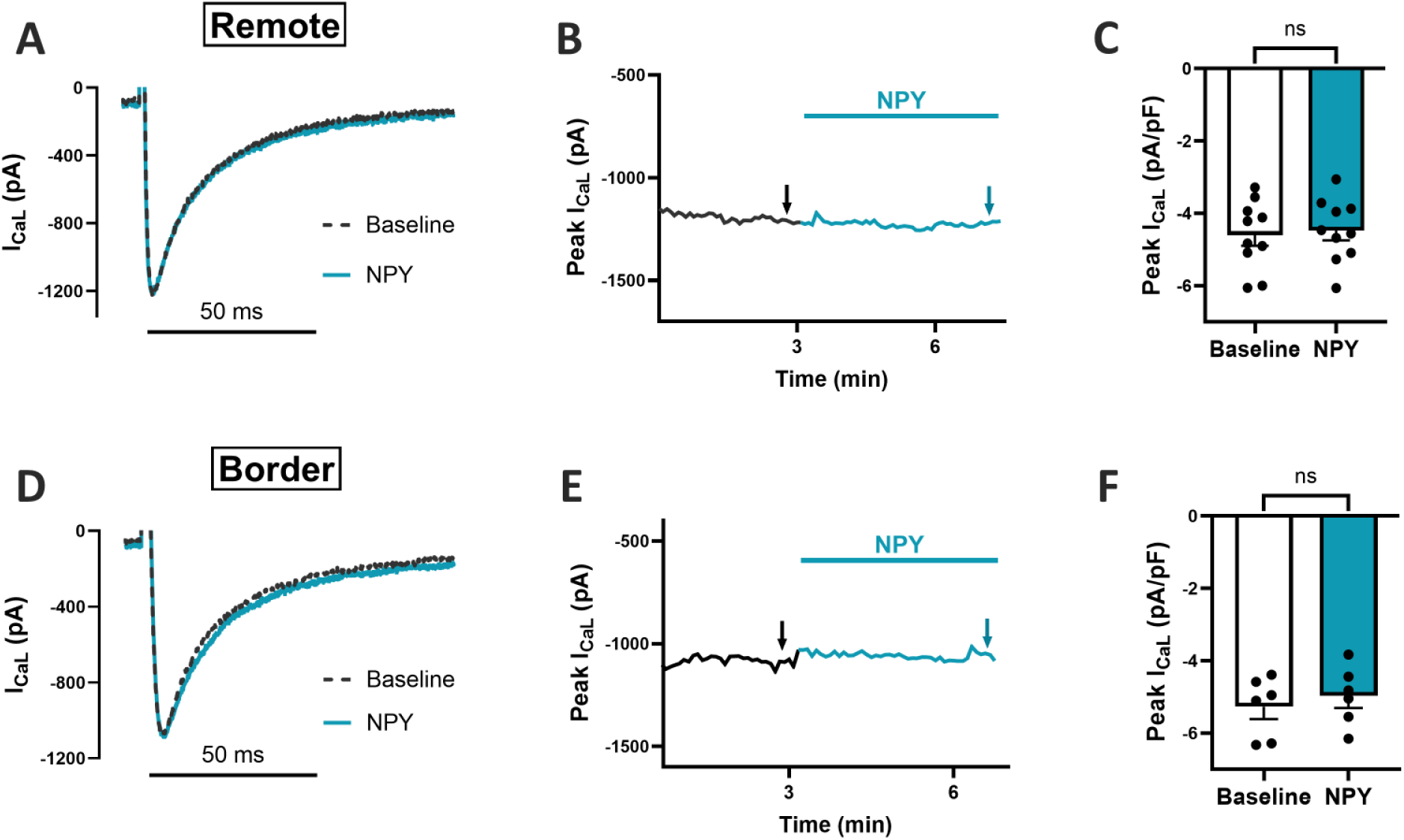
NPY fails to enhance I_CaL_ in remote and border zone ventricular myocytes post-MI. (A) Representative whole-cell I_CaL_ traces recorded under control conditions (dashed black line) and following application of 100 nM NPY (blue line). (B) Time course from an example myocyte showing no change in I_CaL_ amplitude after NPY application. (C) Group data reveal no significant difference (2.27 ± 3.13% decrease) in average I_CaL_ response to NPY (baseline: –4.60 ± 0.30 pA/pF; NPY: –4.48 ± 0.28 pA/pF; p = 0.4387, N/n = 4/10). (D) Representative I_CaL_ traces under baseline conditions (dashed black line) and after 100 nM NPY application (blue line). (E) Time course shows minimal change in I_CaL_ with NPY exposure. (F) Average data indicate a non-significant trend toward reduction (6.76 ± 3.46%) following NPY treatment (baseline: –5.27 ± 0.34 pA/pF; NPY: –4.97 ± 0.33 pA/pF; p = 0.1105, N/n = 4/6).

We then examined the response to NPY in the presence of NE. In myocytes from the remote zone (Figure 5A-C), the baseline current density was 6.1 ± 3.5 pA/pF. This increased to 14.3 ± 0.983 pA/pF following exposure to 30 nM NE. This represents a 132% increase over baseline (p < 0.0001). Following addition of NPY, the current density decreased to 13.5 ± 1.16. However, this effect was not statistically significant (p = 0.10, N/n = 5/10; Figure 5C). In border zone myocytes (Figure 5D-F), the peak Ca^2+^ current density was 5.4 ± 0.32 pA/pF under baseline conditions. This increased to 11.9 ± 0.887 pA/pF following exposure to 30 nM NE. This represents a 118% increase over baseline (p < 0.0001). Following addition of 100 nM NPY, the current density decreased to 10.2 ± 0.834 pA/pF. This represents 25.2% decrease in the response to NE (p = 0.018, N/n = 5/9; Figure 5F). These results indicate that the stimulatory effect observed in response to NPY alone in control myocytes is lost in both remote and border zone cells following MI. However, the inhibitory effect observed in the presence NE can still be observed following MI, especially in border cells.

**Figure 5.**
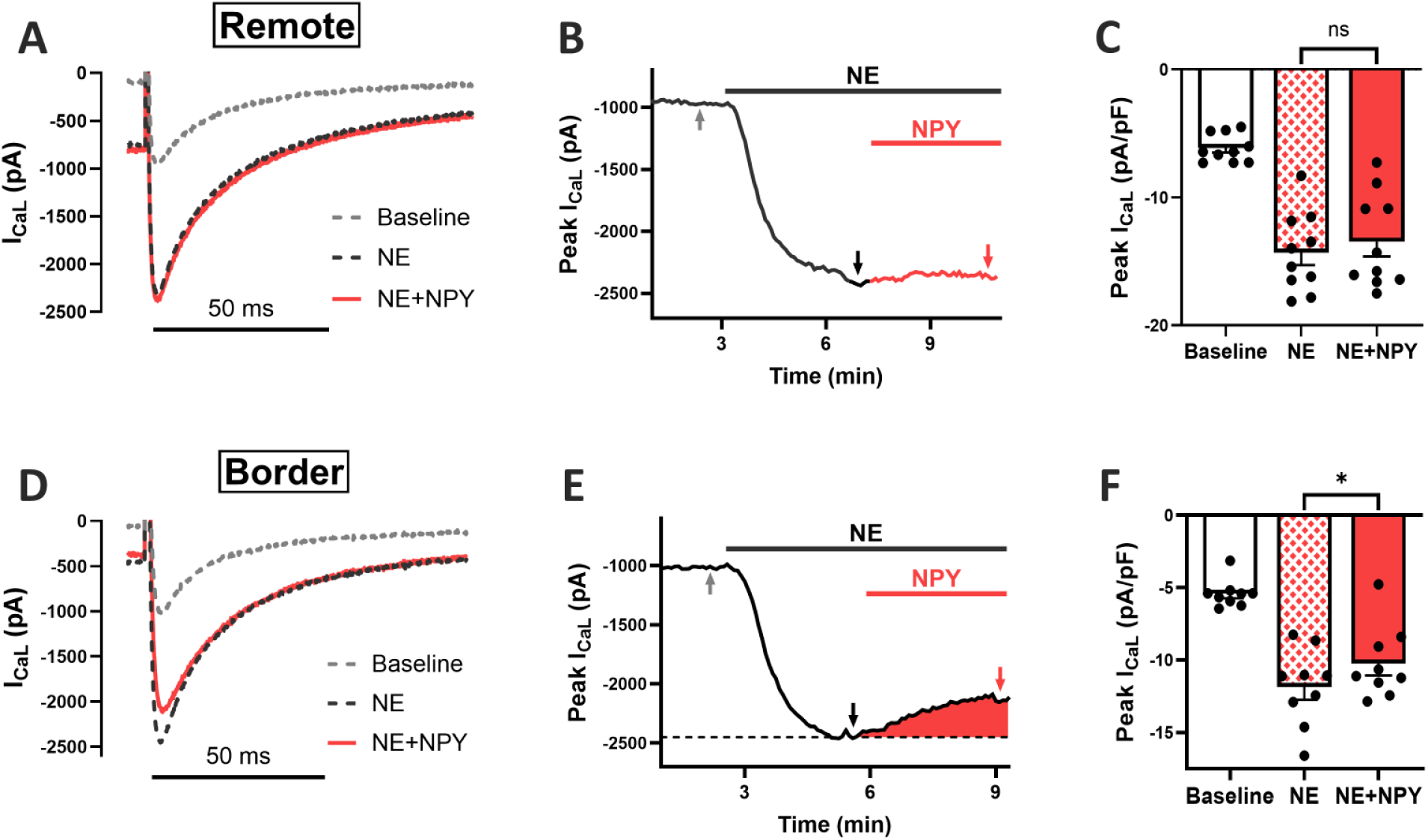
NPY mediated reduction of NE-stimulated I_CaL_ is regionally altered in MI-PVM. (A) Representative I_CaL_ traces recorded under baseline (dashed gray line), 30 nM NE (dashed black line), and NE + 100 nM NPY (red line) conditions. (B) Time course demonstrates a small decrease in peak I_CaL_ with NPY application. (C) Mean current densities show a trend toward reduction by NPY with an average 12.5 ± 7.06% decrease (NE: –14.3 ± 0.98 pA/pF; NE+NPY: –13.5 ± 1.16 pA/pF; p = 0.1032, N/n = 5/10). (D) Representative I_CaL_ traces recorded under baseline conditions (dashed gray line), with 30 nM NE (dashed black line), and after NE + 100 nM NPY application (red line). (E) Time course reveals consistent reduction in I_CaL_ by NPY. (F) Average data confirm a significant decrease of 23.3 ± 7.90% in current amplitude upon addition of NPY (NE: –11.9 ± 0.89 pA/pF; NE+NPY: –10.2 ± 0.83 pA/pF; p = 0.0176, N/n = 5/9).

## Discussion

In the present study, we found that NPY alone stimulates the L-type Ca^2+^ current in ventricular myocytes from normal healthy pigs. Previous studies have looked at the effect on the Ca^2+^ current in myocytes from different small animal models, with varying results. In guinea-pig ventricular myocytes, NPY was reported to actually inhibit the Ca^2+^ current [17], while in rat ventricular myocytes NPY has been reported to have either no effect [15], or to stimulate the Ca^2+^ current [16]. The explanation for the variability in the response is unclear. It is also unclear what the significance of the response to NPY alone is, since in vivo, sympathetic stimulation of the heart would be expected to release NPY together with NE. For that reason, we examined the Ca^2+^ current response following exposure to NE, where we found that NPY actually has an inhibitory effect (see Graphical Abstract).

There has been at least one previous study that has looked at the interaction between the effects of NPY and β-adrenergic stimulation. However, the authors focused primarily on the effects that NPY had on contraction of normal rat ventricular myocytes [16]. It was found that NPY alone enhanced contraction, while in the presence of the β-agonist, isoproterenol, NPY had an inhibitory effect. It was demonstrated that the stimulatory effect of NPY alone could be explained by an increase in the Ca^2+^ current. However, the authors concluded that NPY inhibition of contraction in the presence of isoproterenol was due to an increase in the transient outward K^+^ current and a subsequent shortening of action potential duration. While we did not look at the effects of NPY on contraction in pig myocytes, one might expect a similar overall response. NPY alone would be expected to enhance contractility due to an increase in the Ca^2+^ current, while in the presence of β-adrenergic stimulation it would be expected to inhibit contraction. However, any inhibitory effect is unlikely to be explained by changes in the transient outward K^+^ current, since pig myocytes lack such a current [18]. Instead, it would be more likely that any inhibitory would be due to antagonism of the effect that β-adrenergic stimulation has on the Ca^2+^ current, which was demonstrated here for the first time.

The present study is also the first to examine the specific receptor subtypes involved in mediating the effects of NPY on the Ca^2+^ current in cardiac myocytes. We found that the stimulatory effect was blocked by the Y_1_ antagonist BIBO, but not the Y_2_ antagonist BIIE. Because Y_1_ receptors are associated with Gq signaling pathways, it is likely that the stimulatory effect that NPY has the Ca^2+^ current in pig myocytes is due to activation of phospholipase C and PKC. This is consistent with the effect that NPY has on Ca^2+^ transients in rat ventricular myocytes [15]. It is curious, however, that NPY was reported to have no effect on the Ca^2+^ current in those cells. That could be due to the reported variability in the effect of PKC on the cardiac Ca^2+^ current [20].

The ability of NPY alone to actually inhibit the Ca^2+^ current in guinea-pig myocytes was attributed to the activation of a G_i_-dependent mechanism, presumably involving inhibition of AC [17]. Although it was not determined what receptor subtype was involved, it is likely that Y_2_ receptors were involved, since they are known to signal through G_i_. The fact the stimulatory effect of NPY alone was not enhanced in the presence of BIIE in our study suggests that inhibition of basal AC activity does not affect the Ca^2+^ current in pig ventricular myocytes. However, BIIE did block the inhibitory effect of NPY in the presence of NE, consistent with the idea that the inhibitory effect is due to Gi-dependent inhibition of AC and cAMP production in the presence of β-adrenergic receptor stimulation. Consistent with this conclusion, we observed that NPY also inhibited the cAMP regulated Cl^-^ current activated in the presence NE and this effect was blocked by the Y_2_ antagonist BIIE, but not the Y_1_ antagonist BIBO (Supplementary Figure 1).

The effects of NPY on the Ca^2+^ current recorded in myocytes isolated from normal healthy pigs are important for understanding the role of this neuropeptide under physiological conditions. However, NPY has also been implicated in sympathetic induced changes in electrical activity that contribute to cardiac arrhythmogenesis following MI. Clinically, elevated NPY levels in the coronary sinus correlate with greater arrhythmia burden, larger infarct size, and impaired recovery of ventricular function post-STEMI (ST-elevation myocardial infarction) [12,13,21,22]. A reduction in ventricular fibrillation threshold independent of β-adrenergic stimulation has also been observed in the Langendorff-perfused rat heart following MI [13]. This effect was attributed specifically to the activation of Y_1_ receptors. Furthermore, the upregulation of Y_1_ receptors was found to be proarrhythmic in a mouse model of subarachnoid hemorrhage [23]. Y_1_ receptor mediated increase in Ca^2+^ influx in cardiac myocytes could conceivably contribute to these observations. However, while we did observe a Y_1_ receptor mediated stimulatory effect of NPY alone on the Ca^2+^ current in control myocytes, the net effect of NPY on the Ca^2+^ current in the presence of NE was inhibitory, both in myocytes from normal health animals as well as myocytes isolated from MI hearts.

It is interesting to note that we observed a loss of the stimulatory effect that NPY alone has on the Ca^2+^ current following MI, while the inhibitory effect that occurs in the presence of NE was largely intact. The inhibitory affect did appear to be diminished in myocytes isolated from remote regions of the infarcted heart, while it seemed to remain intact in myocytes from the border zone. It is unclear what these changes might mean in terms of any potential arrhythmogenic mechanism. It could suggest that effects on Ca^2+^ transients independent of the Ca^2+^ current, such as has been reported in the rat [15], may be more important. It could also be that NPY effects on other types of ion channel activity are important. Finally, it may suggest that other mechanisms, such as presynaptic modulation of sympathetic or parasympathetic neurotransmitter release [11] or vasoconstriction due to vascular smooth muscle contraction [24,25], may play a role.

## Acknowledgements

This work was supported by the National Institutes of Health National Heart, Lung, and Blood Institute Grants P01 HL164311, R01 HL161122, and P20 GM130459.

