## Supplemental Data for "Neuropeptide Y regulation of L-type Ca^2+^ channel activity is altered following chronic myocardial infarction"

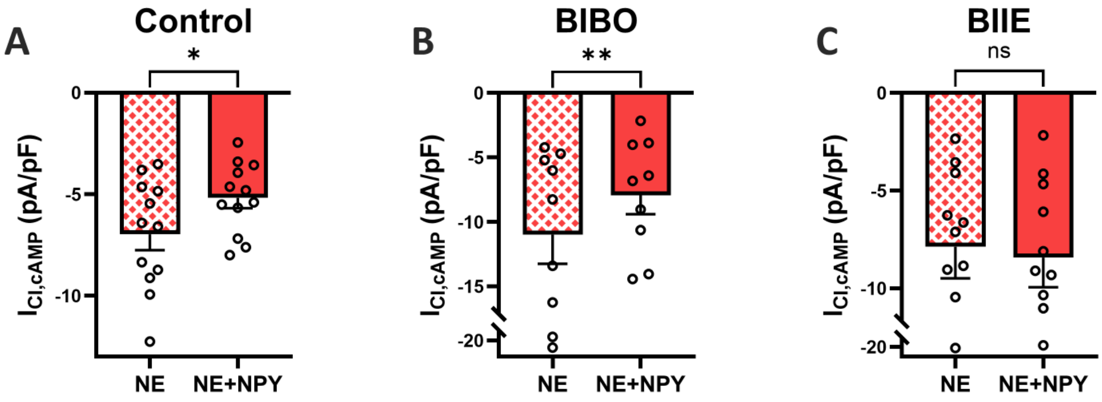


**Supplemental Figure 1. NPY suppresses the cAMP-dependent chloride current in control PVM via $Y_2$ receptor activation.**

(A) NE+NPY co-application significantly reduced $I_{Cl,cAMP}$ relative to NE alone (NE: –6.97 ± 0.78 pA/pF; NE+NPY: –5.18 ± 0.50 pA/pF; 28.0 ± 8.95% decrease; p = 0.0197). (B) This suppression was still present with $Y_1$ receptor blockade (NE: –11.0 ± 2.26 pA/pF; NE+NPY: –7.93 ± 1.48 pA/pF; 26.0 ± 5.27% decrease; p = 0.0088). (C) Blockade of $Y_2$ receptors abolished this suppression (NE: –7.88 ± 1.62 pA/pF; NE+NPY: –8.43 ± 1.52 pA/pF; 10.9 ± 4.78% increase; p = 0.1032).
